# Mathematical modelling of mutant glycoprotein fate in the ER

**DOI:** 10.64898/2025.11.30.691435

**Authors:** Daniele Di Bella, Andrea Lia, Pietro Roversi

**Affiliations:** Institute of Agricultural Biology and Biotechnology (IBBA), National Research Council of Italy (CNR), via Bassini 15, I-20133 Milano, Italy; Department of Environmental Sciences, Informatics and Statistics, Ca’ Foscari University, Dorsoduro 3246, 30123 Venice, Italy; Institute of Nanotechnology (NANOTEC), National Research Council of Italy (CNR), Campus Ecotekne, Via Monteroni, I-73100 Lecce, Italy; Department of Experimental Medicine, University of Salento, Campus Ecotekne, Via Monteroni, I-73100 Lecce, Italy; Leicester Institute of Chemical and Structural Biology and Department of Molecular and Cell Biology, University of Leicester, Henry Wellcome Building, Lancaster Road, Leicester LE1 7HR, England, United Kingdom

**Keywords:** ERQC, ERAD, UGGT, ER lectins, ER mannosidases, misfolded glycoprotein, ODE

## Abstract

**Background:** The endoplasmic reticulum (ER) coordinates glycoprotein quality control through competing pathways: ER quality control (ERQC) and ER-associated degradation (ERAD). The fate of a glycoprotein is tied to its *N*-glycan(s) structure, which serves as a “molecular barcode” for quality control. In particular, modulation of ERQC/ERAD can increase leakage of responsive (*i.e*., non-inactive) glycoprotein mutants that remain misfolded yet retain biological activity once outside the ER. Rescue of secretion of such mutants is a promising therapeutic strategy for the therapy of congenital rare disease due to a missense mutation in a secreted glycoprotein gene, but the relative merits of UGGT *vs*. EDEM inhibition to release a mutant from the ER remain to be explored.

**Methods:** An ordinary differential equation (ODE) framework (“ER Glycoprotein Outcome”, ERGO) evaluates time-dependent ER concentrations of 13 *N*-glycospecies of a single mutant glycoprotein (G1M9, G1M8B, G1M8C, G1M7BC, M9, M8A, M8B, M8C, M7AB, M7AC, M7BC, M6, M5) within the ER lumen, under a set of chosen hypotheses. All reactions follow mass-action kinetics with ER enzymes/lectins kinetic constants and concentrations as fixed parameters. The ODE system is solved in `Python` with adaptive Runge-Kutta integration. Simulations initialise from a single glycoform (M9, 1 *µ*M) and track glycoform distributions over ∼28 h. Starting glycoform concentrations and overall time can be customised. Given the lack of comprehensive kinetic data in the literature, parameter values reflect educated guesses, rather than measured constants.

**Results:** Four simulation scenarios explore the dynamics of ERQC *vs* ERAD: (A) baseline scenario with all enzymes active; (B) UGGT inhibition, which abolishes re-glucosylation, disrupts CNX/CRT cycles, and accelerates substrate flux toward ERAD-prone glycoforms; (C) inhibition of all ER mannosidases, preventing trimming-dependent ERAD commitment and prolonging ER retention of early glycoforms; (D) combined UGGT and ER mannosidases inhibition, mimicking experimental rescue of misfolded substrates by blockade of both glucosylation-driven ER retention and trimming-driven degradation signals. We further quantify the rate of CNX/CRT engagement (‘carousel rides’) by combining analytical estimates with Monte Carlo–style simulations of individual mutant glycoprotein fates.

## Introduction

In the Endoplasmic Reticulum (ER) of Eukaryotes, the fate of a newly synthesised glycoprotein depends on a delicate balance between quality-control mechanisms that promote correct folding and pathways that eliminate mis-folded glycoproteins via ER-associated degradation [1]. The ER surveillance system exploits modifications of *N*- glycans linked to the glycoprotein (see Figure 1) as molecular signposts that determine whether the glycoprotein will dwell in the ER for assisted folding or be funnelled to destruction [2]. Two glycan-modification patterns govern this binary fate decision, with two enzymatic systems dynamically acting on them, creating a kinetic tug-of-war [3, 4]:

(i) **The ER retention pathway:** a mono-glucosylated glycoprotein bearing a terminal glucose on branch A of the glycan (Figure 1) is recognised by ER lectins calnexin and calreticulin (CNX/CRT), which retain it in the ER and hand it over to foldases [5, 6]. This retention is dynamically controlled by two opposing enzymes:
  (a) ER Glucosidase II (GluII) removes the terminal glucose, releasing the glycoprotein from CNX/CRT and allowing it to proceed along the secretory pathway [7, 8];
  (b) UDP-glucose glycoprotein glucosyltransferase (UGGT) – the ERQC folding sensor: it adds back a glucose to *N*-glycans of a misfolded glycoprotein, regenerating a mono-glucosylated glycan that prolongs interaction with CNX/CRT and grants further folding attempts [9, 10, 11]. Notably, UGGT action is not restricted to fully mannosylated glycans: *in vitro* UGGT can reglucosylate Man_9_GlcNAc_2_ (M9) as well as partially trimmed Man_8_GlcNAc_2_ (M8B or M8C^1^) and Man_7_GlcNAc_2_ (M7BC) structures, albeit with decreasing efficiency [12, 13].
(ii) **The ERAD commitment pathway** an ER glycoprotein is progressively demannosylated by ER *α*mannosidases until the Man(*α*1,6)Man moiety on Branch C is recognised by lectins OS-9/XTP3-B – the canonical ERAD receptors – and the misfolded client is delivered to the cytoplasmic degradation machinery [14, 10]. ER *α*-mannosidases remove mannose residues specifically from the M9 core, producing trimmed structures (M8A, M8B, M8C; M7AB, M7BC, M7AC, etc.) [15]. EDEM2 catalyses the first cut from M9 to the specific M8B glycoform [16, 15], followed by EDEM3 [17] (and partly EDEM1 [18, 19, 20, 21]), which removes additional mannoses to generate M7BC, M7AB and smaller glycoforms. Strikingly, purified EDEMs can demannosylate even glucosylated intermediates (G1M9, G1M8 and G1M7) *in vitro*, implying that the presence of a terminal glucose does not inherently block mannose trimming [22, 23].

**Figure 1.**
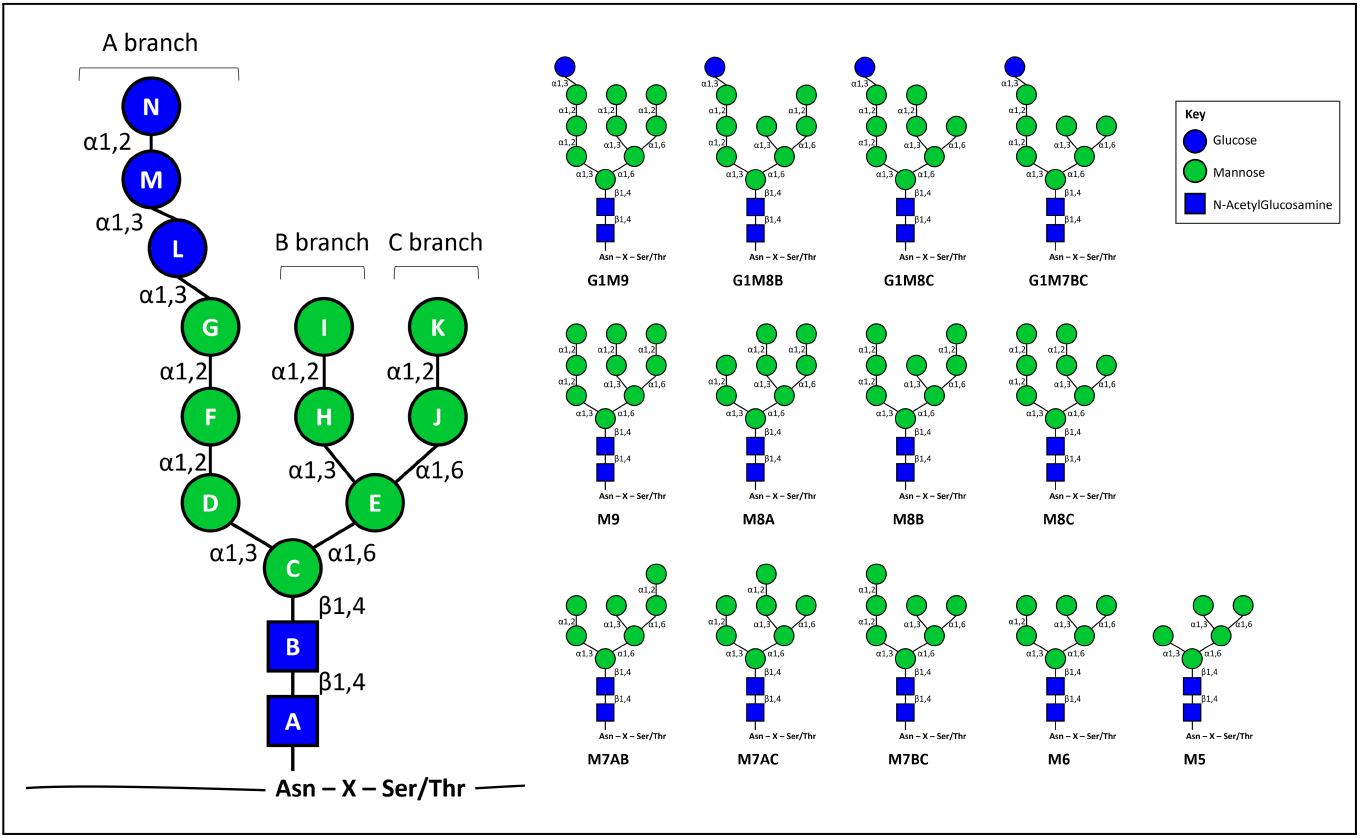
Schematic representation of the *N*-glycan. The image illustrates the canonical branching topology of the Glc_3_Man_9_GlcNAc_2_ structure and the 13 glycoforms mentioned in this study. Branches A, B, and C are indicated following standard glycan nomenclature, with each monosaccharide depicted according to [40]. Blue squares denote GlcNAc residues, blue circles represent glucose units on the A-branch, and green circles correspond to mannose residues. *N*-glycosylation motif: Asn-X-Ser/Thr, where X is not Pro. This representation provides a visual reference for the *N*-glycan structural context underlying analyses in this study.

Crucially, the ability of UGGT to act on partially trimmed glycans and the one of EDEMs to trim glucosylated substrates means that a glycoprotein can simultaneously bear both retention and degradation signals. The ultimate fate of each misfolded/not yet folded glycoprotein therefore emerges from a kinetic competition between UGGT, which continually restores the ER retention signal, and EDEMs, which progressively mark the substrate for degradation. Once a C-branch Man(*α*1,6)Man moiety) is exposed, the misfolded glycoprotein binds to the lectins OS-9/XTP3-B and crosses an irreversible commitment threshold — the “event horizon” — beyond which re-glucosylation can no longer rescue it (Figure 2)^2^.

**Figure 2.**
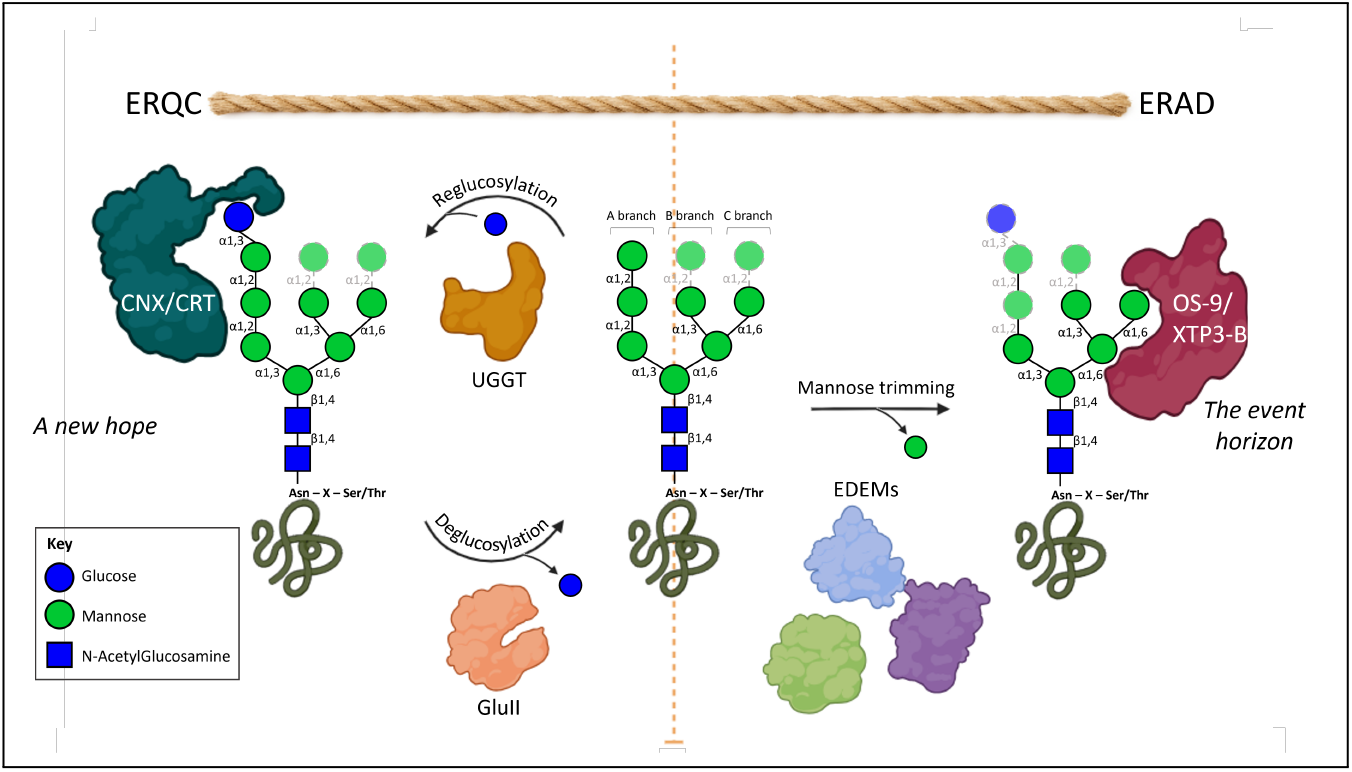
Tug-of-war between ERQC, representing the “new hope” of folding and secretion, and ERAD, representing the “event horizon” for a misfolded glycoprotein. Misfolded glycoproteins can be reglucosylated by UGGT and re-engage calnexin/calreticulin (CNX/CRT, left) or undergo mannose trimming that exposes the C-branch signal recognised by OS-9/XTP3-B (right), irreversibly committing them to ERAD. The *N*-linked glycans are represented with the standard shapes and colours for monosaccharides in glycans [40]: blue squares: N-acetyl glucosamine (GlcNAc); green circles: mannose (Man); blue circles: glucose (Glc). Solid (transparent) sugar residues are (not) necessary for lectin recognition and binding. Some elements of the figure were created with BioRender.com

While our qualitative understanding of these processes has been strengthened by recent biochemical and structural work, fundamental questions remain open: how is the misfolded glycoprotein’s fate (ER retention *vs* degradation) decided? What factors determine the outcome of this kinetic tug-of-war? Are misfolded glycoprotein fates dictated primarily by the concentrations of ERQC and ERAD enzymes and lectins, or do the relative rates of re-glucosylation, deglucosylation, mannose trimming, and escape to the Golgi play an equally decisive role? How many re-glucosylation cycles can a misfolded glycoprotein undergo, before being captured by OS-9/XTP3-B and crossing the event horizon?

Answering these questions requires a quantitative framework that integrates the known biochemistry of ER quality control into formal time-dependent models. Such a framework enables modelling of the ERQC/ERAD system as a set of simultaneous competing reactions – UGGT-driven retention *vs* mannosidase-driven degradation – with enzymes’ and lectins’ concentrations and association/dissociation on/off rates of ERQC/ERAD enzymes and lectins as tunable parameters.

We describe here the first implementation of a system of ordinary differential equations (ODEs) named ERGO – ER Glycoprotein Outcome, also evoking the Latin *ergo* (“therefore”), to emphasise that mutant glycoprotein fate results from the balance of ER retention, escape to the Golgi, and ERAD. ERGO simulates the time-dependent concentrations of glycoforms under diverse ERQC/ERAD model assumptions and formalises the “mannose-timer” hypothesis of ER quality control [26]: newly synthesised glycoproteins enter the calnexin (CNX) folding cycle, are subject to slow mannose trimming, and only after trimming (*i*.*e*., loss of critical mannoses) become committed to ERAD. Figure 1 illustrates the glycoforms relevant to the present work. In the model, CNX-bound glycoforms do not spontaneously convert into a folded species; instead, their fate depends on the competition between retention*/*re-glucosylation (via UGGT) and demannosylation (via EDEMs*/*ER EM or analogous mannosidases). The “clock” is effectively the slow mannose-trimming kinetics: proteins that fail to fold (or are folding-resistant mutants) are gradually trimmed and thereby tagged for degradation, while those that would fold before sufficient trimming (if a folding pathway were present) could escape.

In ERGO’s current form, focussed as it is on mutant glycoproteins intrinsically incapable of reaching the native state, no folding-rate term is implemented, and mannose trimming inexorably commits glycoproteins to ERAD or leakage — embodying the minimal “mannose-timer” model. Accordingly, the present ODE formulation of `ERGO` does not include any transition from CNX-bound or monoglucosylated species to a folded, secretion-competent glycoprotein. All glycoforms remain ERQC*/*ERAD clients throughout. Consequently, the model quantifies modulation of the kinetic competition between UGGT-mediated retention and EDEM-mediated ERAD commitment for glycoprotein mutants that never attain their native conformation. Under this assumption, the secretion (SEC) sink represents leakage of misfolded but potentially therapeutically responsive mutants – rather than *bona fide* secretion.

By simulating how mutant glycoprotein populations redistribute among different *N*-glycan structures over time, ERGO enables the role of each kinetic parameter in influencing substrate fate, including the magnitude and direction of its effect, and prediction of the outcomes of enzyme inhibition. Experimental validation of such models would ideally come from time-resolved measurements of glycoform distributions in the ER, obtained through reconstituted *in vitro* systems under defined conditions. Comparing simulated and observed dynamics would allow iterative refinement of model parameters and discrimination among alternative mechanistic hypotheses. Ultimately, this interplay between quantitative modelling and targeted experiments will reveal the main factors that irreversibly commit a misfolded glycoprotein to degradation, clarifying how the complex web of glycan modifications either grants further time for folding or pushes substrates past the “event horizon” towards ERAD.

## Methods

### Model formulation

ERGO provides a quantitative framework for investigating the fate of mutant glycoproteins in the endoplasmic reticulum by formalising the interplay between folding surveillance mechanisms and the pathways that commit substrates to ERAD. All reactions in our system are modelled under the assumption of mass–action kinetics, which states that the rate of a reaction is proportional to the product of the concentrations of the reactants raised to powers corresponding to their stoichiometries. This principle provides the foundation for representing enzymatic transformations, lectin binding, and other ERQC*/*ERAD processes in a unified mathematical framework.

Examples of such reactions treated under mass–action kinetics include:

(i) UGGT-mediated re-glucosylation of an incompletely folded species, *e*.*g*.: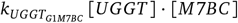, where the *N*-glycan M7BC, with molar concentration [*M* 7*BC*], is converted to G1M7BC by UGGT, with molar concentration [*UGGT*]; the subscript on *k* identifies the enzyme performing the reaction and the resulting glycoform (see the next section for the choice of formats for kinetic constant labels);
(ii) removal of the innermost glucose on an *N*-glycan branch A by GluII, *e*.*g*.: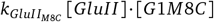.;
(iii) progressive mannose trimming by ERManI and EDEM1–3, represented similarly, by a rate constant multiplied by the corresponding enzyme and substrate concentrations;
(iv) endomannosidase (ER EM)-mediated cleavage of a Glc-Man disaccharide from branch A in monoglucosylated species (G1M9, G1M8B, G1M8C, G1M7BC) to yield downstream glycoforms (M8A, M7AB, M7AC, M6), each step encoded by a *k*-term specifying the resulting glycoform and multiplied by [*ER EM*] · [*S*_*j*_], where [*S*_*j*_] indicates the concentration of the *j*-th glycoform;
(v) mutually exclusive binding equilibria between glycoforms and lectins CNX or OS-9, captured by rate constants whose subscripts identify the type of lectin interaction (binding, on, or unbinding, off) and whose corresponding mass–action terms take the form [*CNX*] · [*S*_*j*_] or [*OS*9] · [*S*_*j*_].

For an enzymatic step where an enzyme *E* acts on a glycoform *S*_*j*_ (reducing its molar concentration [*S*_*j*_]) to produce another glycoform *S*_*k*_ (thus increasing [*S*_*k*_]), mass–action kinetics gives

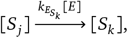

with rate law

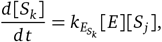

and the corresponding depletion of the substrate

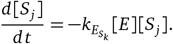

Similarly, according to mass-action kinetics, binding interactions between glycoforms and lectins follow standard bimolecular association and unimolecular dissociation kinetics. For instance, CNX binding to a free glycoform *S*_*j*_ is represented as

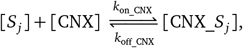

With

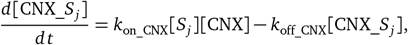

And

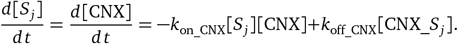

Since total lectin pools are finite,

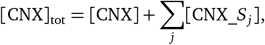

and the free lectin concentration is

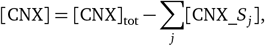

introducing nonlinearity in the system through terms of the form

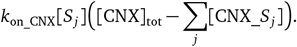

On the other hand, enzymatic conversions remain linear under the assumption of constant enzyme pools^3^.

Collectively, these contributions lead to the general differential equation governing the concentration of the general glycoform *S*_*j*_,

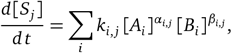

which aggregates all processes producing or consuming *S*_*j*_. Each term in the sum corresponds to a specific reaction *i*, with positive or negative rate constant *k*_*i*,*j*_, involving enzymes or lectins *A*_*i*_ and glycoforms *B*_*i*_ raised to their respective stoichiometric exponents *α*_*i, j*_ and *β*_*i, j*_. The ensemble of differential equations encompassing all *j* glycoforms represents the interconnected biochemical steps that define the structure of the ERQC and ERAD network. An example of such a complete system of ODEs corresponding to a certain set of molecular hypotheses is provided in Supplementary Sections 2.1 and 2.2.

Integrating this system yields time-resolved predictions of how the mutant glycoprotein population redistributes among three global fates – ER retention, escape to the Golgi, and degradation – under specified combinations of UGGT activity, mannosidase availability, and lectin occupancy. These simulations reveal how reglucosylation and demannosylation function as complementary regulatory arms: the former sustaining substrate recovery within the folding cycle, the latter driving commitment towards disposal.

As with any mathematical abstraction, the model is defined by explicit assumptions including, in our case:

- a single *N*-glycan per substrate and absence of tri- or di-glucosylated states (mimicking an ALG6(-/-) background);
- a translation-inhibited regime in which total glycoprotein mass is conserved (analogous to cycloheximide experiments), ensuring no influx from synthesis;
- in absence of knowledge about relative affinities of CNX *vs* CRT and OS-9 *vs* XTP3-B, and with all glycospecies belonging to a single glycoprotein, one monoglucosylated glycan-binding lectin only (“CNX”) and one ERAD-type glycan-binding lectin only (“OS-9”);
- unsaturated, constant enzyme and lectin pools, with no cooperativity or Michaelis–Menten saturation;
- finite, mutually exclusive CNX/OS-9 binding enforcing realistic competition for lectin availability;
- absence of any folding terms, so the model represents glycoproteins that never reach the native state; CNX association only causes ER retention, it is not conducive to attainment of fold. This is because ERGO is meant to support the study of pathology-associated glycoprotein mutants which are misfolded and retained, but functionally responsive once escaped from the ER;
- unidirectional export of free species to the Golgi, and irreversible ERAD of OS-9–bound forms; as the model does not include any folded state or folding-promotion step, all glycoforms remain ERQC/ERAD clients. Thus, the SEC sink represents leakage rather than *bona fide* secretion;
- a single rate of escape to the Golgi and a single rate of ERAD after binding OS9 lectin, irrespective of free or OS9-bound glycospecies, respectively [27].

Within these assumptions, the model enables to explore how specific enzymatic or lectin-driven perturbations reposition the balance between ER retention, escape to the Golgi, and degradation; to derive insights into the thresholds that switch a substrate from ERQC to ERAD; and to quantify how changes in UGGT activity, mannosidase flux, or lectin saturation reshape the landscape of mutant glycoprotein fate decisions. Because the framework captures only the core regulatory logic of ERQC/ERAD, it provides a foundation that can be systematically refined through the insertion of experimental data, and expanded by incorporating additional ER factors.

As already mentioned, ERGO lacks any term that converts a CNX-associated monoglucosylated species into a folded, secretion-competent glycoprotein. As visible in the ODEs (*e*.*g*., G1M9, G1M8B, G1M8C, G1M7BC equations in Supplementary Section 2.1) there is no transition from any glycoform to a “folded” state; UGGT reglucosylates, enabling renewed CNX binding, but this interaction does not lead to productive folding. CNX association is modelled purely as reversible retention, with no downstream folding outcome. All glycoforms, including G1M9/G1M8/G1M7BC, remain ERQC/ERAD clients until demannosylation commits them to ERAD (Supplementary Section 2.2). Therefore, ERGO quantitatively represents mutants that never fold, consistent with a focus on unstable or misfolded pathological variants.

### Choice of parameters and initial and boundary conditions

As mentioned in the paragraphs above, the ERQC/ERAD ODE model depend on parameters such as ERQC/ERAD enzymes’ and lectins’ concentrations, enzymes’ rate constants and lectin complexes’ k_on_ and k_off_ rates, and overall rates of mutant glycoprotein escape to the Golgi and degradation. The dependence of the ER fate of a substrate glycospecies on these choices can be explored by running ERGO calculations differing at least by the value of one of the parameters.

Some enzymes can catalyse multiple reactions, and the kinetic constants associated with each reaction are, in principle, reaction-specific. This specificity arises because the kinetic parameters depend on the detailed molecular interaction between enzyme and substrate – an interaction unique to each enzyme:substrate pair – including the orientation of the substrate within the active site, *e*.*g*. the two cleavages performed by GluII differ for their products and rates [8]. To make explicit which binding mode a given rate constant refers to, and to emphasize that a glycospecies may be processed by the same enzyme at different sugar positions with sugar-specific kinetics (e.g., EDEM1 or EDEM3 demannosylation of species retaining intact A or C branches), we adopt the notation `k_<enzyme>_<product>` in our code and the corresponding notation 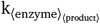 in this text ^4^. Similarly, to represent lectins binding and unbinding rate constants, we adopt, respectively, the notation `kon_<lectin>` and `koff_<lectin>` in our code, and *k*_⟨on_ lectin⟩_ and *k*_⟨off_ lectin⟩_ in our text.

ERGO will be most useful when experimental measurements of these parameters will be available – *e*.*g*. from a series of *in vitro* kinetic experiments, each using one ERQC/ERAD enzyme/lectin and a purified glycoform. In absence of such experimental values, we made the assumptions listed in Table 1. This enables us to illustrate what use can be made of ERGO calculations in the limiting scenario of a full kinetic characterisation of the relevant enzymatic and lectin complex association/dissociation reactions.

**Table 1.**
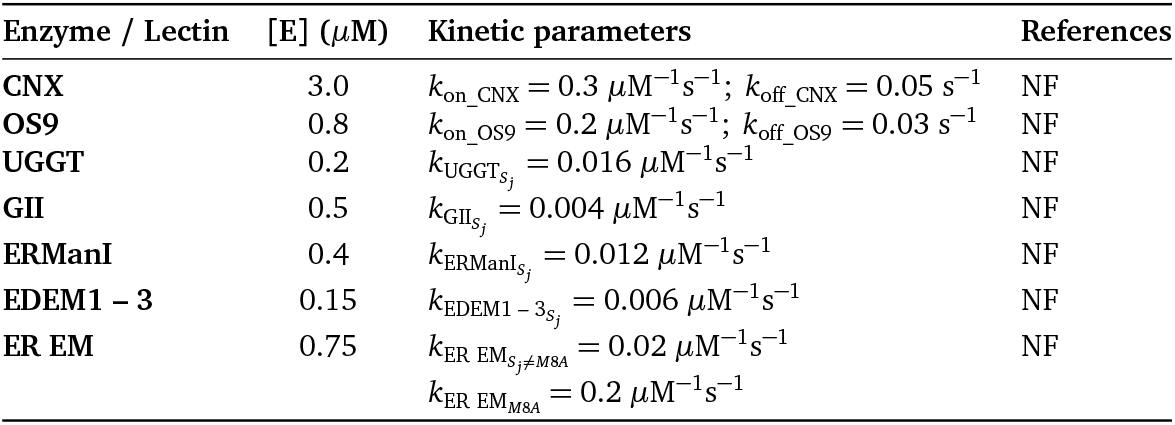
Kinetic parameters of the ER enzymes and lectins implemented in the Python code of our model. The notation 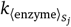 follows the convention 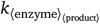 introduced in the section “Choice of parameters and initial and boundary conditions” : it represents the mass-action rate constant at which the ⟨enzyme⟩ produces the glycoform *S*_*j*_. Similarly, *k*_on__ ⟨_lectin_⟩ and *k*_off__ ⟨_lectin_⟩ represent, respectively, the binding and unbinding rates of the specified ⟨lectin ⟩. We searched the literature for each parameter: “NF” in the References column means that no reliable source was found. All values (including enzyme concentrations) are reported exactly as implemented in the code.

Simulations are performed under the assumption of *ALG6*^(–/–)^ background, in which the only monoglucosylated *N*-glycans are those produced by UGGT: ERGO currently has no provision for OST glycoprotein synthesis, ER Glu I, malectin or the first ER Glu II-mediated cleavage. The initial total mutant glycoprotein mass is set to 1.0 *µ*M, distributed entirely as a mutant glycoprotein bearing a non-glucosylated nine-mannose *N*-glycan (“M9”). Secreted mutant glycoproteins are accumulated in a single “SEC” compartment that serves as an irreversible sink representing leakage to the Golgi and beyond. A single degradation sink term takes care of modelling ERAD degradation of glycospecies associated to the OS-9 lectin [27]. At each point in time, the overall concentration of ERAD-degraded glycospecies is computed as the complement of the sum of the ones of glycoforms remaining in the ER plus those in the SEC pool, with respect to the initial total concentration of mutant glycoprotein.

The single rate of escape to the Golgi for all free glycospecies and the single rate of ERAD for all OS9-bound glycospecies have been set to values of the order of 10^−3^*s*^−1^. Unimolecular kinetic constants have been chosen to be in the order of magnitude of 10^−3^ – 10^−2^*s*^−1^. CNX and OS-9 total pools are fixed at 3.0 *µ*M and 0.8 *µ*M, respectively. The total and constant concentrations of ER enzymes and lectins are reported in Table 1.

### Numerical integration

The ODE system is integrated using the `solve_ivp` function from `SciPy`, with adaptive time-stepping and relative and absolute tolerances of 10^−6^ and 10^−9^, respectively. The time span ranges from 0 s to 10^5^ s, and output points are sampled logarithmically to capture early- and late-time dynamics. The integration scheme automatically adjusts step size to maintain numerical stability and accuracy across the different kinetic regimes.

### Post-processing and analysis

The array of integrated numerical ODE solutions **y**(*t*) contains the time-dependent values of concentrations of ER glycospecies and lectin complexes. From this array one can compute the total concentrations of:

- CNX-bound mutant glycoproteins (ER-retained fraction);
- OS-9-bound mutant glycoproteins and ERAD-committed fraction;
- free glycoforms in the ER lumen (potentially secreted fraction); and
- accumulated leaked-to-Golgi pool (SEC).

Simulating four perturbation scenarios, plots of concentration versus time were generated using the matplotlib library. For each scenario, both the aggregate fates and individual glycoform concentrations were visualized to facilitate comparison among scenarios, (and with experimental data once/if available).

## 1 Results

### ODE simulations of mutant glycoprotein fate under four regulatory scenarios

The ERGO system of ODEs quantitatively reproduces the redistribution of a mutant glycoprotein population among the three major fates within the endoplasmic reticulum (ER): retention, escape to the Golgi, and degradation. Four regulatory scenarios were simulated by setting specific catalytic constants to zero in the model described in Supplementary Sections 2.1 and 2.2:

- (A) Uninhibited system — baseline parameters;
- (B) UGGT inhibited — all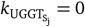;
- (C) Mannosidases inhibited — all 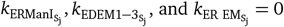;
- (D) Combined inhibition — all UGGT and mannosidases catalytic constants set to zero.

In the uninhibited baseline condition (Figure 3A), UGGT-mediated re-glucosylation and ER mannosidase-driven demannosylation dictate the time-dependence of glycospecies concentrations. A fraction of mutant glycoproteins (dark blue in the left-hand side panels in Figure 3) sticks in the ER, free or engaged with CNX/CRT or OS-9, while the rest is progressively committed either to escape to the Golgi (gold in the left-hand side panels of Figure 3) or ERAD (grey in the left-hand side panels in Figure 3). The heatmaps in the right-hand side panels of Figure 3 highlight the concentrations of individual high-mannose glycoforms, with glucosylated intermediates transiently accumulating before disappearing due to either escape to the Golgi or degradation.

**Figure 3.**
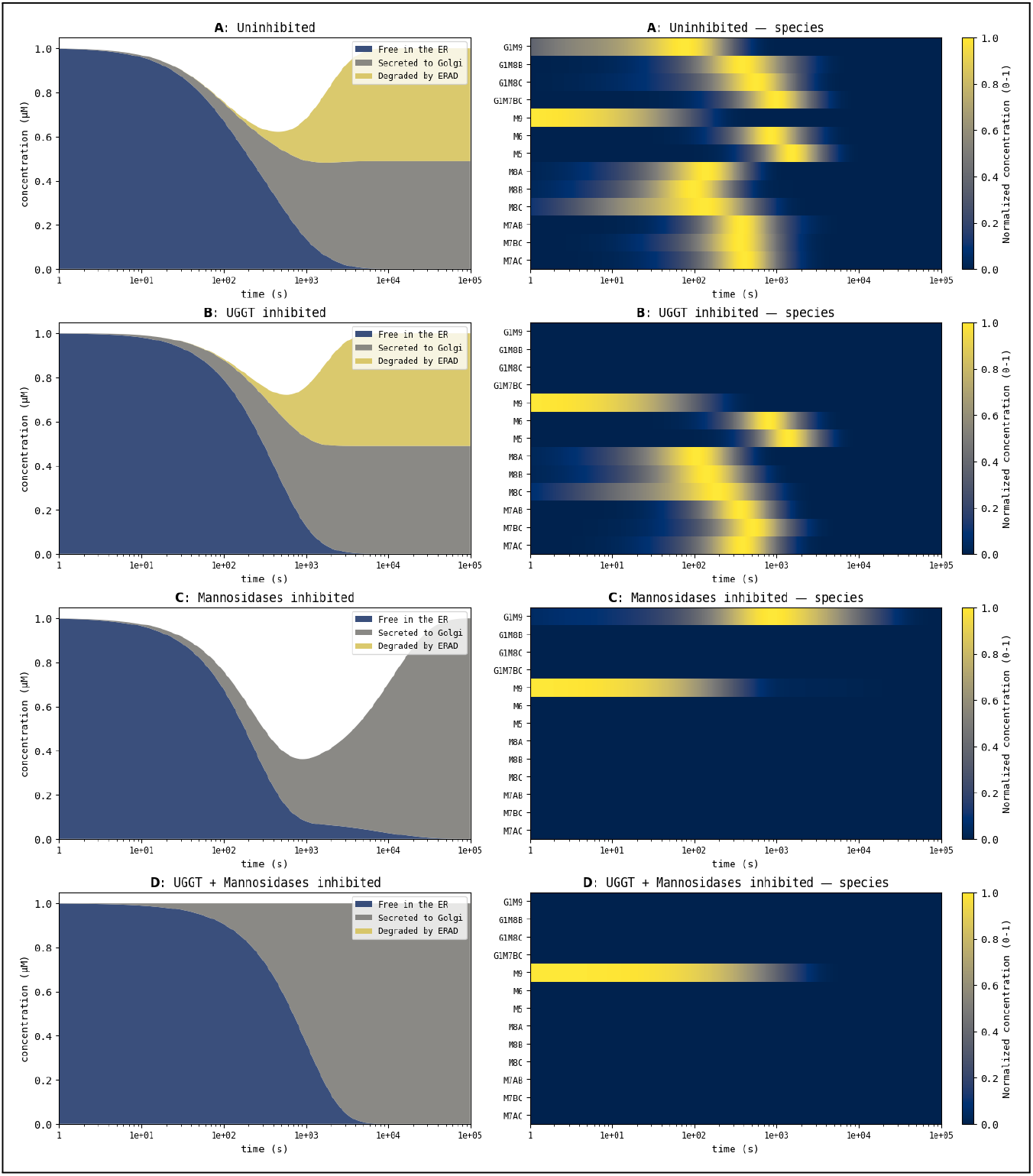
Dynamics of mutant glycoproteins fate under four scenarios of the ERQC*/*ERAD system. For each condition—(A) baseline, (B) UGGT inhibition, (C) mannosidases inhibition, and (D) combined inhibition—the left-hand side panel shows the temporal redistribution of the mutant glycoprotein population among three global fates (free/retained in the ER, escaped to the Golgi, and degraded via ERAD), displayed as a stacked area plot on a logarithmic time axis. Conversely, the right-hand side panel presents heat maps of the concentrations of the 13 free glycoforms, normalized row-wise to highlight their relative temporal dynamics. In the baseline scenario (A), UGGT-mediated reglucosylation and ER mannosidase activity generate a dynamic balance between lectin-bound retention, secretion, and ERAD commitment. UGGT inhibition (B) suppresses re-glucosylation, causing complete absence of glucosylated intermediates, thus preventing access to CNX, and accelerating leakage to the Golgi and ERAD. Mannosidase inhibition (C) blocks demannosylation-dependent ERAD, prolongs ER residence, increases secretion, and stabilizes high-mannose species (notably M9). Combined inhibition (D) produces a kinetically inert system in which the only species is M9.

When UGGT activity is inhibited (Figure 3B), reglucosylation is abolished. This leads to a reduced fraction of mutant glycoproteins interacting with CNX/CRT, reduced ER retention time, and to accelerated secretion/degradation via OS-9–mediated ERAD. The total population shows faster depletion of free mutant glycoproteins and increased degradation, while heatmaps reveal the expected absence of glucosylated species.

The inhibition of ER mannosidases (Figure 3C) prevents the trimming of mannose residues required for ERAD. Escape to the Golgi happens at a faster rate. The ERAD fraction is non-existent, and, coherently with mannosidase inhibition, the heatmaps show longer survival of the M9 and a late peak of the G1M9 glycoform. The increased SEC flux corresponds to increased escape because ERAD commitment is prevented but folding cannot occur.

Simultaneous inhibition of both UGGT and ER mannosidases (Figure 3D) generates a kinetically inert system. The only glycoform is the unprocessed M9 species, with null engagement in ERAD and prolonged ER retention, eventually progressing towards leakage to the Golgi. UGGT and ER mannosidase joint inhibition abolishes the dynamic ERQC/ERAD turnover observed in the uninhibited scenario, boiling down to M9 escape to the Golgi over the timescale examined.

### ODE-guided identification of discriminant experimental observables

Overall, ERGO simulations capture the impact on mutant glycoprotein fate of modulating ERQC and/or ERAD enzymes. The calculations can be repeated to test different ERQC/ERAD models or choosing different concentrations and rate constants, to predict how UGGT and ER mannosidases differentially influence the partitioning of mutant glycoproteins between ER retention, leakage to Golgi, and degradation, linking enzyme activity to both the kinetics of individual glycoforms and the overall distribution of the mutant glycoprotein pool. Thus, ERGO provides a framework to explore mechanistic ERQC/ERAD assumptions. By selectively modifying hypotheses — *e*.*g*., including or excluding ER endomannosidase (ER EM), or permitting simultaneous association with both calnexin and OS-9 — one can predict which glycoform will respond most strongly in their temporal concentration profiles. The same holds for different choices of ERQC/ERAD component concentrations and rate constants. In presence of experimental observations for all or some of the time-dependent glycospecies concentrations (from *in vitro* simplified systems or from *in cellula* data) ERGO can be used to assess the relative likelihood of such ERQC/ERAD models.

Glycospecies exhibiting the largest changes in concentration when comparing alternative ERQC/ERAD models, constitute desirable experimental targets, because their time-dependent behaviour is most sensitive to the underlying molecular details. Measuring the time-dependence of their concentrations *in vitro*, for instance by mass spectrometry or lectin-affinity assays [28], would provide maximal information to discriminate among competing mechanistic hypotheses. The ERGO framework therefore also constitutes an *in silico* tool aiding experimental design, highlighting where measurement effort will most effectively assist evaluation of the relative likelihood of ERQC/ERAD molecular models.

Last but not least, ERGO enables fine quantitative interrogation of ERQC/ERAD processes. In addition to linking individual glycospecies to their macroscopic fates, the framework can be leveraged to resolve finer kinetic features that are otherwise experimentally elusive. As we illustrate in the next section, this includes estimating the frequency of specific microscopic events—such as the number of CNX/CRT binding cycles per second (“carousel rides”) experienced by a misfolded glycoprotein — thereby extending the utility of ERGO from experimental design to the mechanistic dissection of ERQC/ERAD phenomena.

### Quantifying the number of CNX/CRT “carousel rides”: two complementary strategies

In the context of ERQC models, each binding event of a monoglucosylated glycoform to the lectin CNX may be thought of as one “ride” on a molecular carousel: a round on the quality-control merry-go-round before either successful folding (and onward secretion), further processing, or disposal via OS-9-mediated ERAD. For each specific ERQC/ERAD model, ERGO can address the question: on average, how many times per second does a misfolded glycoprotein engage in a CNX association event?

To answer this question, two strategies are available:

1. a deterministic ODE model to compute the instantaneous binding rate of CNX to monoglucosylated species and convert it into a per-molecule average rate over time. We refer to this approach as the “analytical estimation” of the rate of misfolded glycoprotein CNX association;
2. a stochastic, Monte Carlo–style simulation of individual misfolded glycoprotein trajectories through the ERQC/ERAD network, counting each CNX binding event as a discrete “carousel ride” and estimating the empirical average rate from many simulated trajectories. We refer to this approach as the “Monte Carlo-style estimation” of the rate of mutant glycoprotein CNX association.

The two approaches yield orthogonal population-level and molecule-level estimations of misfolded glycoprotein CNX engagement frequency, enriching our mechanistic insight into the retention–folding–disposal decision circuit.

### Average CNX-association rate: analytical estimation

Firstly, we define the set of monoglucosylated glycospecies:

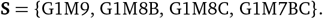

Then, we define the molar instantaneous *total* association rate of free CNX to this pool, *i*.*e*., the total micromoles of CNX complexes formed by misfolded glycoproteins in **S**, at time *t*, per mole of CNX per volume:

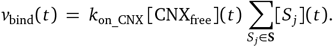

where:

- *k*_on_CNX_ is the calnexin association rate constant with units *µ*M^−1^ s^−1^.
- [CNX_free_](*t*) is the concentration of unbound CNX molecules at time *t* (*µ*M).
- [*S*_*j*_](*t*) for *S*_*j*_ ∈ **S** is the concentration of misfolded glycoprotein molecules belonging to the general CNX-binding glycospecies *S*_*j*_ at time *t*. This is also expressed in *µ*M.

The product 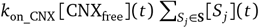 therefore has units (*µ*M^−1^ s^−1^) · *µ*M · *µ*M = *µ*M s^−1^.

To obtain the per-glycoprotein instantaneous binding rate (that is, the expected number of CNX association events experienced by a single glycoprotein molecule per unit time), we have to divide the total instantaneous association rate, *v*_bind_(*t*), by the total concentration of glycoproteins. Formally, let **A** be the set of all glycoforms, and

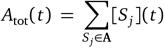

the total concentration of misfolded glycoproteins at time *t*. The instantaneous per-glycoprotein rate is then

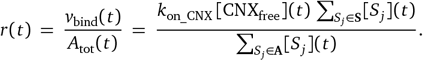

The numerator has units *µ*M s^−1^, and the denominator has units *µ*M. Thus, *r*(*t*) has units *µ*M s^−1^*/µ*M = s^−1^, coherently with it being a per-molecule rate.

The time-averaged per-glycoprotein association rate 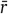 is the average of *r*(*t*) over a certain time T:

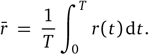

Substituting *r*(*t*) = *v*_bind_(*t*)*/A*_tot_(*t*) gives

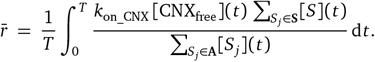

For the mutant glycoprotein and the ERQC/ERAD model described in Supplementary Sections 2.1 and 2.2, initialised with the parameters reported in Table 1 and averaging over a time T=28 h, one obtains: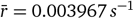.

### Average CNX-association rate: Monte Carlo-style estimation

The same time-averaged per-glycoprotein CNX association rate 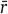 can also be estimated by averaging the count of CNX association events over a large number of Monte Carlo simulation of *N* individual misfolded glycoprotein molecules following stochastic trajectories. At any one time *t*, each glycoprotein molecule can occupy one of *j* discrete states (*S*_*j*_) – which can be thought of as nodes of a graph. For a mutant glycoprotein, states include free glycoforms, CNX-bound species, OS9-bound species, as well as states indicating that the molecule has been committed to escape to the Golgi (`SEC`) or degradation (`DEG`). The graph in Figure 4 represents the mutant glycoprotein and the ERQC/ERAD system implemented in the ODEs in Supplementary Sections 2.1 and 2.2, with transition probabilities (see below) at time t=0.

**Figure 4.**
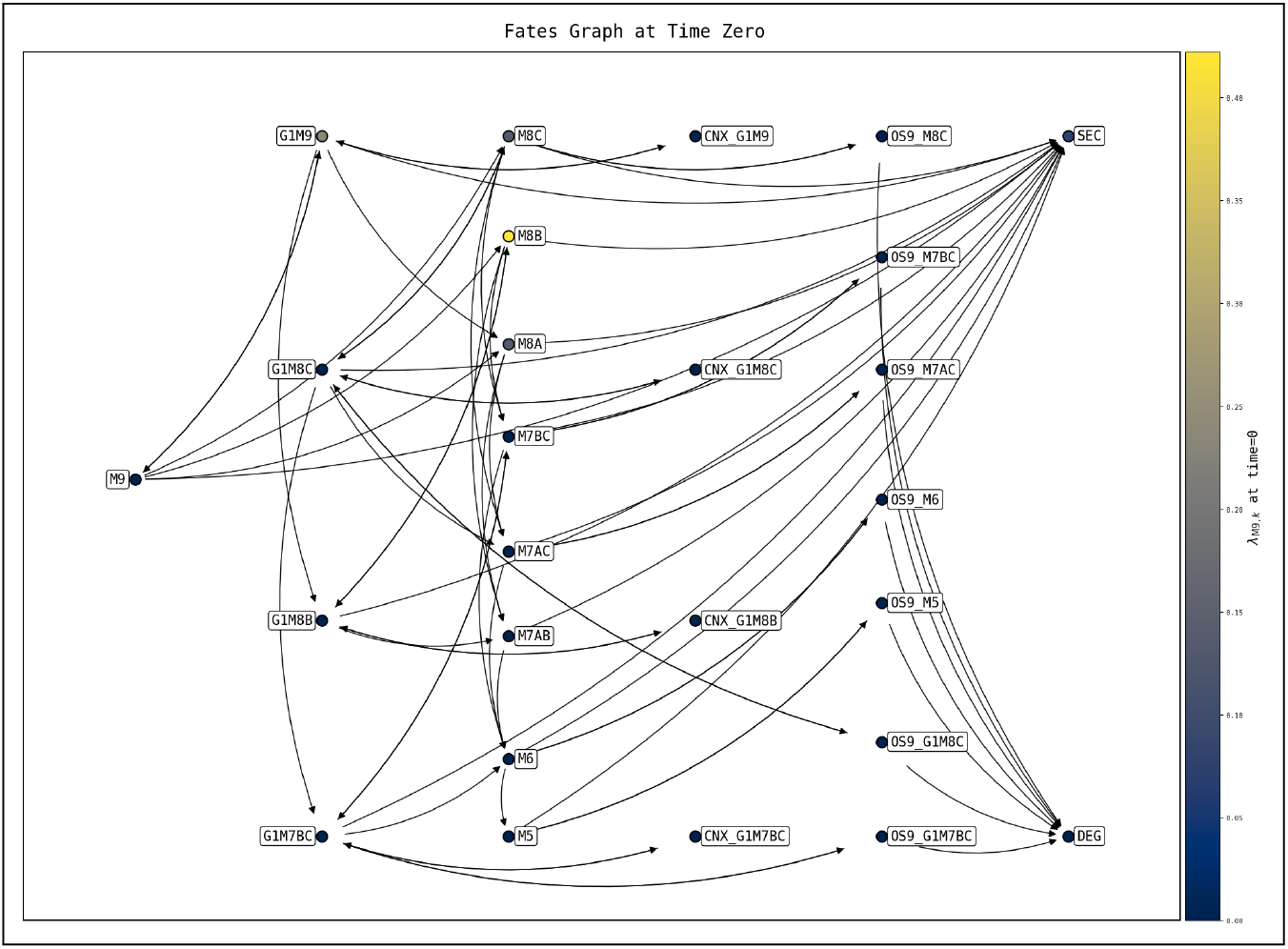
Graph representing an ERQC*/*ERAD system at time=0. The graph refers to the mutant glycoprotein and the ERQC/ERAD model described in Supplementary Sections 2.1 and 2.2, initialised with the parameters reported in Table 1, with transition probabilities at time t=0. Nodes represent free glycoforms (e.g. `M9`, `G1M9`, `M8A` …), lectin-bound complexes (prefixed `CNX_` or `OS9_`), and sink states (`SEC`, `DEG`). Edges denote transitions among states (enzymatic conversions, lectin binding/unbinding, secretion, ERAD routing). Node fill colour encodes the one-step reachability probability from the start state `M9` at *t* = 0, *i*.*e*. for each target node *k* the summed outgoing rate from `M9` towards that target (*λ*_M9,*k*_, *t* = 0), divided by the total outgoing rate (*Λ*_*M* 9_, *t* = 0).

The implemented Monte Carlo algorithm advances time according to the discrete time grid defined by the ODE integrator: *t*_0_, …, *t*_*N*_ with step *Δt*_*i*_ = [*t*_*i*_; *t*_*i*+1_) on each interval [*t*_*i*_, *t*_*i*+1_). The *N* trajectories (each describing the transformations undergone by one individual glycoprotein) are modelled as Markov processes, meaning that what will happen in the small time interval *Δt*_*i*_ = [*t*_*i*_; *t*_*i*+1_) depends only on the extension of *Δt*_*i*_ and on the state *S*_*j*_ in which the tracked misfolded glycoprotein is at time *t*_*i*_.

We start by defining *Λ*_*j*_ (*t*_*i*_), the “total hazard rate” from state *S*_*j*_:

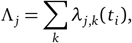

where *λ*_*j*,*k*_ (*t*_*i*_) is the “hazard rate”, computed at time *t*_*i*_, at which a misfolded glycoprotein moves from state *S*_*j*_ to a neighbouring, reachable state *S*_*k*_. In other words, *λ*_*j*,*k*_ (*t*_*i*_) is the instantaneous probability per unit time, expressed in *s*^−1^, at which our tracked misfolded glycoprotein goes from *S*_*j*_ to *S*_*k*_. According to the mass-action framework we outlined above, we define

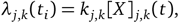

where:

- *k*_*j*,*k*_ is the enzymatic rate constant or the lectin associations/disassociation rate for the transition from state *S*_*j*_ to state *S*_*k*_;
- [*X*]_*j*,*k*_(*t*_*i*_) is the concentration at time *t*_*i*_ of the lectin or enzyme involved in the same transition: this is the reason why *λ*_*j*,*k*_ varies with time.

*Λ*_*j*_ (*t*_*i*_) gives the chance of a transition from state *S*_*j*_ to any of its reachable states at *t*_*i*_. Thus, 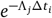can be taken as the probability that no transition from state *S*_*j*_ happens during *Δt*_*i*_. The probability for the tracked misfolded glycoprotein to undergo any transition from state *S*_*j*_ in the same time interval can then be computed as:

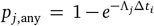

To estimate the average per-glycoprotein CNX association rate, we simulate *N* individual trajectories for a fixed maximum time *T* ^(*tr*)^; some of these trajectories can terminate in either escape to the Golgi or degradation. We call 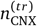 the number of CNX binding events (*S*_*j*_ → *CNX* _*S*_*j*_ transitions) experienced during the simulated lifetime *T* ^(*tr*)^ of the trajectory *t r*. The code detects these events by checking that a transition occurs from a free state *S*_*j*_ to a target state whose name begins with the prefix `“CNX_”`, and increments a per-trajectory counter when such transitions occur. This approach allows us to estimate the Monte Carlo estimate of the mean number of CNX-association events per glycoprotein per second 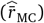:

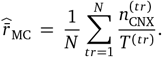

For the mutant glycoprotein and the ERQC/ERAD model described in Supplementary Sections 2.1 and 2.2, initialised with the parameters reported in Table 1, over a time T=28 h, and setting *N* = 2000, the Monte Carlo estimator has the value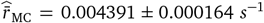. Figure 5 shows that, as *N*→ ∞,, 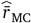 converges to the value 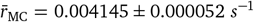 (ergodicity)^5^, which is close to the analytical value for the same model 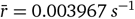.

**Figure 5.**
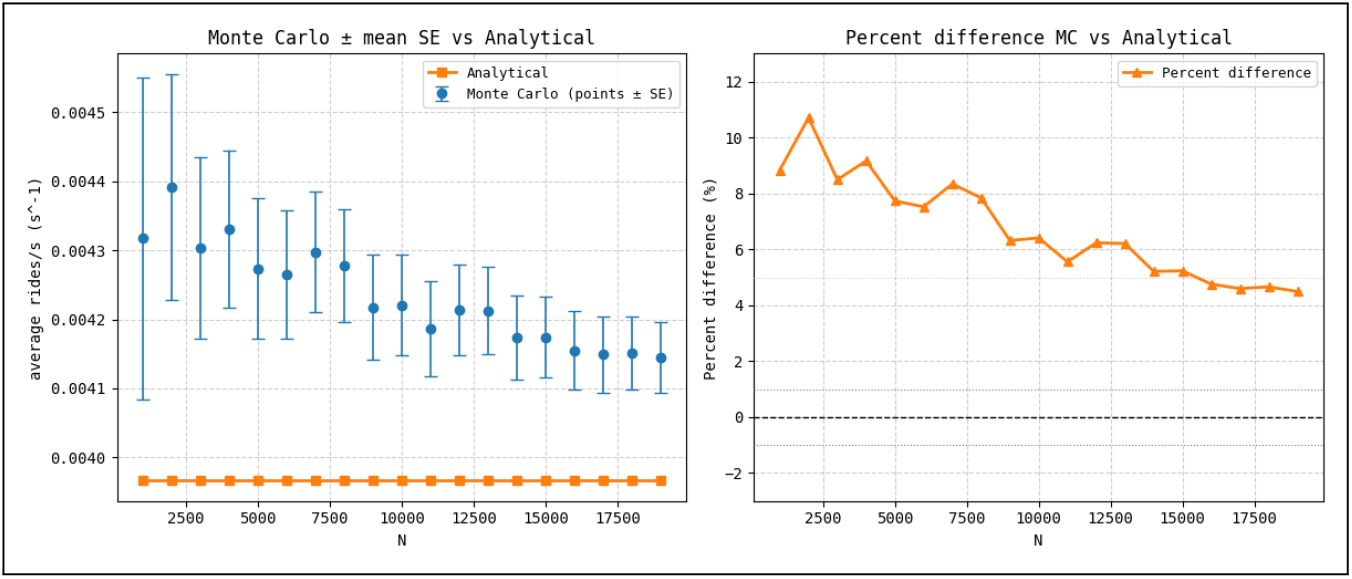
Comparison between Monte Carlo-style and analytical estimates of the mean number of carousel (CNX) rides per second. Left-hand side panel: Monte Carlo estimates computed for different numbers of trajectories (*N*) of the ERQC/ERAD system in Figure 4. Each 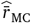 is shown with its standard error (SE) obtained as 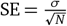. The analytical value (orange squares and connecting line) is constant across all simulations, as expected from the model definition, and reported for comparison purposes. Right-hand side panel: percent difference between the Monte Carlo estimations and the analytical value 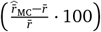, as *N* increases.

## 2 Discussion

ERGO is a quantitative framework for *in silico* modelling of the fates of a mutant glycoprotein molecule subject to ERQC and ERAD within the endoplasmic reticulum. The stochastic (single-molecule) version of the model using kinetic Monte Carlo simulations reproduces population heterogeneity, where individual misfolded molecules follow distinct trajectories of ER retention, escape to the Golgi or degradation. Parameter sensitivity and robustness can be evaluated over the Monte Carlo trajectories.

ERGO depends crucially on ERQC/ERAD enzyme and lectins concentrations and rate constants, and overall rates of escape to the Golgi and degradation. For the ERGO modelling to be useful, these parameters need reliable estimates, such as the ones measurable for a given glycoprotein *in vitro* by mixing a pure glycoform with the ERQC/ERAD components known to interact with it and following the time-dependence of the substrate and product glycospecies by HPLC, lectin binding, or fluorogenic assays.

This first *in silico* reconstitution of the ERQC/ERAD system is intentionally minimalist, isolating the core reactions that determine *N*-linked glycoproteins’ fate. In later phases, this system can be made incrementally more complex, *e*.*g*. by removing the hypothesis of ALG6(-/-) background and introducing rate of glycoprotein synthesis by OST, and/or ER GluI activity and the first glucose cleavage by ER GluII – which would grant initial rounds of CNX association via the OST-transferred glucosylated glycan – or modelling the activity of CNX/CRT-associated foldases aiding transition from misfolded to folded states.

The ERGO *in silico* framework can be used to test alternative hypotheses regarding the molecular mechanisms of ERQC/ERAD. *e*.*g*., different models, all using the same experimentally measured parameters but differing for the molecular events described by the equations, will predict their own “event horizon” in terms of the average time beyond which escape to the Golgi drops below a given threshold. The model whose prediction comes closer to the experimentally measured observation will have the highest likelihood. Discrepancies between predictions and observations will guide modification of the models, *e*.*g*. by addition of ERQC/ERAD components, modelling of multi-component complexes, or enforcement of substrate-specificity on certain components.

Mutant glycoproteins are associated with pathogenic conditions (*e*.*g*. lysosomal storage diseases [29, 30, 31], fibrotic conditions [32, 33] or other congenital rare disease [34]). UGGT or EDEM modulation alters the balance between ER retention and ERAD for misfolded, pathology-associated glycoprotein mutants, thereby controlling their leakage from the ER. For responsive mutants that retain activity despite misfolding, promoting leakage can be a route to functional rescue [35, 36, 37]. The most likely ERGO models will serve as predictive tools to explore ERQC component modulation (*e*.*g*., UGGT or EDEM inhibition/knockdown), recapitulating phenotypes observed in cells and congenital glycosylation disorders. Of course, for ERGO predictions to be accurate in cellular biology and medical contexts, determination of concentrations and rate constants will need *in cellula* measurements. The workflow and kinetic parameters established on test case single-glycan mutant glycoproteins will need generalising to other/poly-glycosylated ones. Measurements conducted in cells from congenital rare disease patients carrying a responsive glycoprotein mutant will be important tests of whether ERGO predictions can reproduce the observed kinetic competition between ER-retention and ERAD that characterises pathology. Once adequate models using reliable parameter estimates are obtained, the ERGO framework will inform on the relative merits of modulation of UGGT *vs* EDEM in rescue of secretion therapy for the treatment of congenital rare disease due to responsive glycoprotein mutants (overexpression, knockdown, or chemical inhibition [35, 36, 38, 39, 37]).

If ERGO is to be used to study wild-type glycoprotein fates, it will need extending by explicit incorporation of a productive folding pathway, which is absent in the current implementation. At present, CNX-associated glycospecies are retained in the ER and can only return to their free state or undergo further demannosylation; no term promotes them to a folded, secretion-competent form. A natural refinement will be therefore to introduce a folding term that removes CNX-bound monoglucosylated species from the ERQC pool and transfers them to a new “folded” species class contributing to a *bona fide* folded, secreted pool, distinct from the present leakage sink. Mathematically, this could mirror the existing ERAD sink: just as OS-9–bound glycoforms feed the *k*_*ERAD*_ degradation term, each *CNX* _*S*_*j*_ complex would feed a single folding rate constant *k*_FOLD_ that converts *CNX* _→*S*_*j*_ *FOLD* (→ *SEC*). This would allow ERGO to model both misfolding-prone mutants (where *k*_FOLD_ = 0) and partially foldable, pathology-associated mutants whose export reflects competition among CNX retention, productive folding, and ERAD commitment. Incorporating such a folding channel would significantly broaden ERGO’s applicability to physiologically relevant and therapeutic scenarios.

## Conclusion

In this work we developed an *in silico* quantitative framework for analysing the fates of a mutant glycoprotein molecule within the endoplasmic reticulum. We model the ERQC/ERAD system through an ODE system, and then use it to simulate the potential fates of mutant glycoproteins in the ER under four different experimental conditions. The same ODE system gives quantitative insight on the average number of CNX association events per unit time, either by analytical estimation or by a stochastic Monte Carlo method. The comparison between analytical predictions and Monte Carlo simulations show that our model formalises soundly the ERQC/ERAD system, establishing a platform for future integrations and extension.

## Data and code availability

The source code for the modelling work described in this manuscript is publicly available from the `GitHub` repository: https://github.com/Daniele-Di-Bella/ERGO. The repository includes the `Python` code as a `Google Colab notebook`. The code is released under the MIT licence. Please cite the code as: Di Bella D (2025)`ERGO` [code]. GitHub. https://github.com/Daniele-Di-Bella/ERGO.

## Author contributions

D.D.B., P.R., A.L.: Conceptualization, experimental design, and writing.

## Competing interests

The authors declare no financial, personal, or professional competing interests.

## Grant information

P.R. was the recipient of a LISCB Wellcome Trust ISSF award, grant reference (2018-2020) 204801/Z/16/Z and a Wellcome Trust Seed Award in Science, (2019-2020) grant reference 214090/Z/18/Z. D.D.B. is supported by the Telethon Grant GJC22077 (01/09/2023 – 31/08/2025, to P.R. and Marco Trerotola) for studying the dependence on UGGT1 of the secretion of TDark glycoprotein mutants. A.L. is supported by grant AR.029.2024 NANOTEC LECCE “National Center for Gene Therapy and Drugs based on RNA Technology” EU funding within PNRR CN00000041 SPOKE 10.

## Acknowledgements

We are grateful to Angelo Santino, Antonio Galeone and Carlos P. Modenutti for helpful discussions.

## Supplementary Material

### 2.1 Modelling glycospecies dynamics

1. G1M9, the mono-glucosylated glycoform is – in the *ALG6* KO background – produced only by UGGT-mediated re-glucosylation of M9. Its free concentration is reduced both by ER GluII deglucosylation and by reversible binding to CNX. Because CNX is assumed to display identical association and dissociation kinetics toward all mono-glucosylated glycospecies [41], a single dissociation constant governs the equilibrium of each CNX–glycan complex:

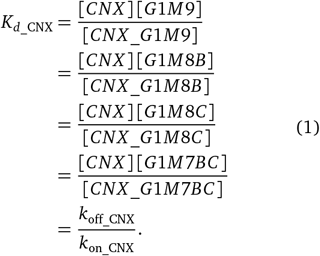

The total CNX concentration therefore reads:

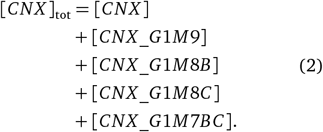

ER ManI, the EDEMs and ER EM can digest G1M9 [24]. UGGT-mediated re-glucosylation slows ERAD, and CNX binding reduces the G1M9 free pool [42, 43]. Collecting all contributions, the time evolution of G1M9 is:

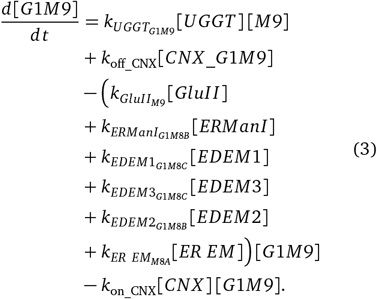
2. G1M8B is generated by ERManI or EDEM2 acting on G1M9, or by UGGT acting on M8B, and it is digested by GluII. The ER endo-mannosidase digests G1M8B to give M7AB [24]. Binding/dissociating to/from CNX also subtracts/adds free G1M8B from the solution:

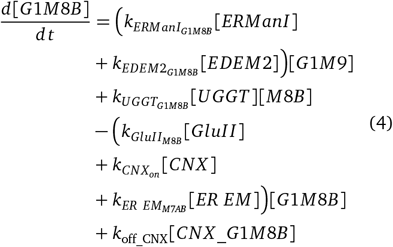
3. G1M8C is generated by EDEM1 or EDEM3 acting on G1M9, or by UGGT acting on M8C, it can be digested by EDEM2 and ERManI and by the ER EM. G1M8C can bind to and dissociate from CNX and OS9. For the latter equilibrium, we assume only one kinetic constant for all glycospecies for the binding to OS9. We write the dissociation equilibrium constant as *K*_*d*_OS9_ with *k*_off_OS9_ and *k*_on_OS9_ rates:

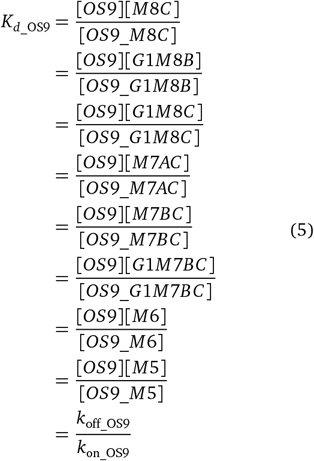

with:

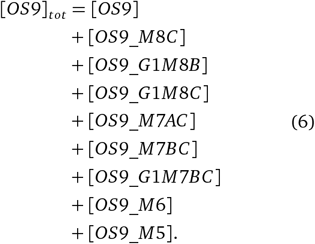

In other words we assume that OS9 has the same on/off kinetics with all its ligand glycospecies and therefore the dissociation constant is one and the same for each OS9 complex with any of its ligand glycospecies. The ER endo-mannosidase (ER EM) digests G1M8C to give M7AC [24]. Binding to CNX also subtracts free G1M8C from the solution. Overall, the rate of change of G1M8C concentration is:

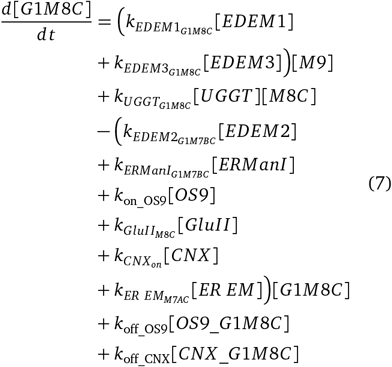
4. M9, the high-mannose glycan, is the default *N*- glycan attached to the nascent mutant glycoprotein in the ALG6 KO background. It is the substrate of UGGT-mediated re-glucosylation [43], and of ER- ManI, EDEM2 and EDEM1/3 demannosylation [42]. It can be generated back when GluII removes the terminal glucose from the mono-glucosylated G1M9 species previously produced by UGGT:

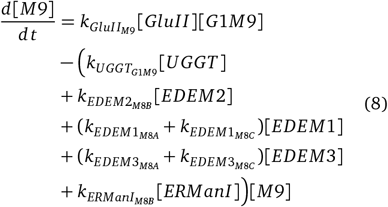
5. M8A, in an ALG6 KO background is generated by: (a) ER EM, from monoglucosylated species G1M9, G1M8B, G1M8C [24]; (b) ER exo-mannosidases acting on M9: not ER ManI or EDEM2 (which are specific for branch B), but EDEM1 and EDEM3[44, 45, 43]. The rate of change in the concentration of M8A can be calculated as:

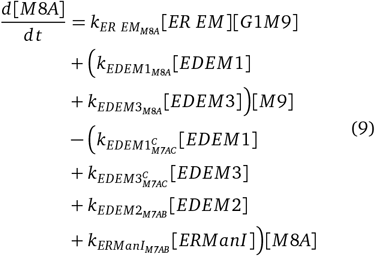
6. M8B is generated by either ERManI or EDEM2 acting on M9, it is digested by EDEM1, EDEM3, and it is reglucosylated by UGGT:

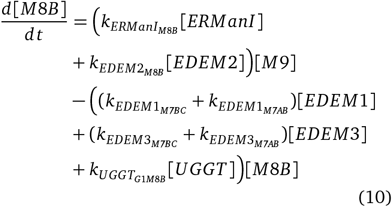
7. M8C is generated by either EDEM1 or EDEM3 acting on M9, or ER GluII acting on G1M8C and it is digested by EDEM2 or ER ERManI, reglucosylated by UGGT and it enters an equilibrium with OS9. The rate of change of free M8C concentration is:

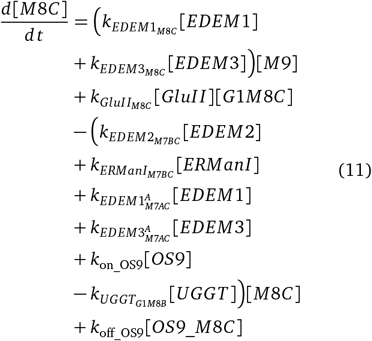
8. M7AB is generated by Man IB or EDEM2 acting on M8A, or by the ER EM endomannosidase complex [24] acting on G1M8B, and it is digested by EDEM1 or EDEM3:

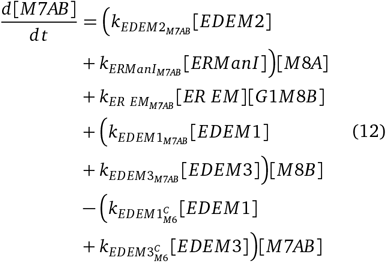
9. M7BC is generated by Man IB or EDEM2 acting on M8C or by EDEM1 or EDEM3 on M8B, it is reglucosylated by UGGT and it is involved in the equilibrium binding to OS9:

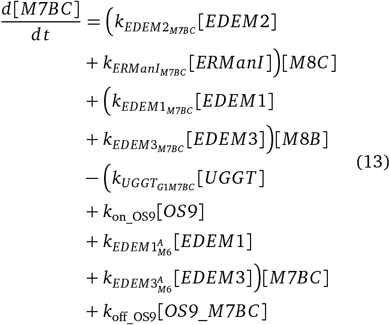
10. G1M7BC is generated by UGGT acting on M7BC, Man IB or EDEM2 acting on G1M8C or by EDEM1 or EDEM3 on G1M8B, it is deglucosylated by ER GluII and it is involved in the equilibria binding to CNX and OS9:

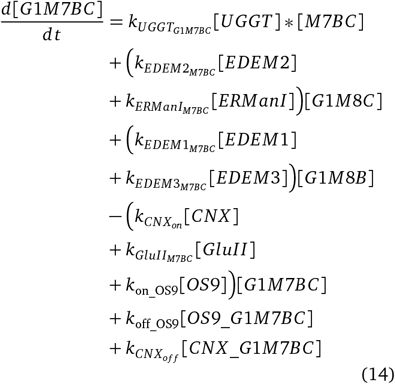
11. M7AC is generated by EDEM1 or EDEM3 acting on M8A or M8C, by the ER EM endomannosidase complex [24] acting on G1M8C, it is digested by EDEM2 or ERManI and it is involved in the equilibrium binding to OS9:

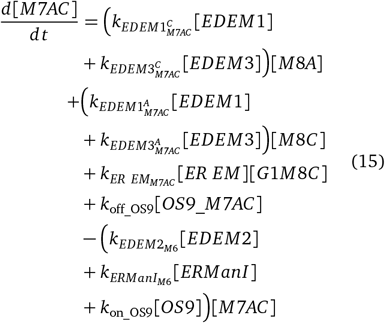
12. M6 is generated by EDEM1,3 acting on M7BC or M7AB, by the ER EM endomannosidase complex [24] acting on G1M7BC, and by ERManI or EDEM2 acting on M7AC; it is involved in the equilibrium binding to OS9:

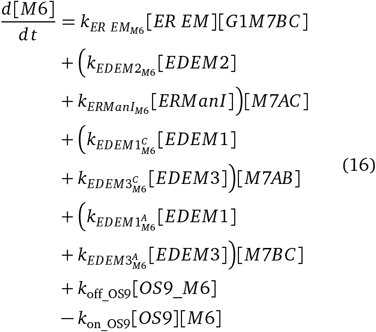
13. M5 is generated by EDEM1,3 acting on M6, and it is involved in the equilibrium binding to OS9:

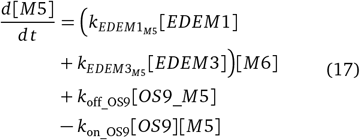

### 2.2 Modelling ERAD dynamics

The sink ERAD term is proportional to the concentration of the glycospecies:OS9 complex, and the K_ERAD_ sink rate is one and the same for all seven OS9 complexes. ER GluII does not remove Glc from glucosylated OS9-bound substrates: as we said, CNX- and OS9-binding are mutually exclusive (a ternary complex of any glycospecies with both lectins is not envisaged).

1. 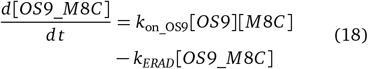
2. 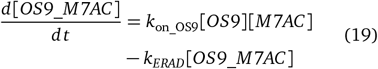
3. 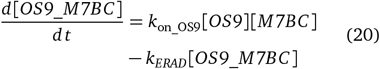
4. 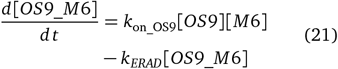
5. 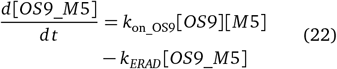
6. 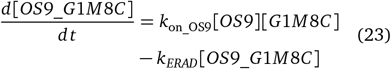
7. 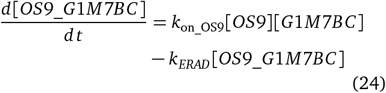

## Footnotes

1 In the nomenclature of this paper, the letter (A, B, or C) indicates the glycan arm from which a mannose residue is removed (see Figure 1), allowing a shorthand description of partially trimmed isomers.

2 Recently, ER Endo-mannosidase activity has been associated with a 800 kDa complex called ER EM [24]. Although the actual Endomannosidase component of the complex has not been discovered, ER EM activity removes the Glc-Man disaccharide from Glc1Man_9–7_GlcNAc_2_ species to yield Man_8–6_GlcNAc_2_ species. This removes the glycoprotein from ERQC by cleaving the mannose on branch A, the one that UGGT re-glucosylates. A candidate enzyme for this activity is ERManIA - for a long time considered a Golgi enzyme until it was found in ER-derived Quality Control Vesicles [25].

3 Our equations are not valid to study the ER fate of an ERQC/ERAD enzyme/lectin when the latter is itself a glycoprotein subject to ERQC/ERAD.

4 The only instances in which this notation would give rise to ambiguities are the symbols for the rates at which EDEM1 and EDEM3 cleave M8A or M8C to give M7AC, and the ones for the rates at which the same enzymes cleave M7AB or M7BC to give M6. The ERGO code has a customisable variable for each of these rates.

5 0.000164 and 0.000052 are, respectively, the standard error associated to 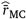 and the standard error associated to 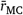.

## Notes

https://github.com/Daniele-Di-Bella/ERGO

### Competing Interest Statement

The authors have declared no competing interest.

